# Foliar pathogens are unlikely to stabilize coexistence of competing species in a California grassland

**DOI:** 10.1101/226928

**Authors:** Erin R. Spear, Erin A. Mordecai

**Affiliations:** Biology Department, Stanford University, Stanford, CA 94305

## Abstract

Pathogen infection is common in wild plants and animals, and may regulate their populations. If pathogens have narrow host ranges and increase with the density of their favored hosts, they may promote host species diversity by suppressing common species to the benefit of rare species. Yet, because many pathogens infect multiple co-occurring hosts, they may not strongly respond to the relative abundance of a single host species. Are natural communities dominated by specialized pathogens that respond to the relative abundance of a specific host or by pathogens with broad host ranges and limited responses to the relative abundance of single host? The answer determines the potential for pathogens to promote host coexistence, as often hypothesized, or to have negligible or even negative effects on host coexistence. We lack a systematic understanding of the impacts, identities, and host ranges of pathogens in natural communities. Here we characterize a community of foliar fungal pathogens and evaluate their host specificity and fitness impacts in a California grassland community of native and exotic species. We found that most of the commonly isolated fungal pathogens were multi-host, with intermediate to low specialization. The amount of pathogen damage each host experienced was independent of host species local relative abundance. Despite pathogen sharing among the host species, fungal communities slightly differed in composition across host species. Plants with high pathogen damage tended to have lower seed production but the relationship was weak, suggesting limited fitness impacts. Moreover, seed production was not dependent on the local relative abundance of each plant species, suggesting that stabilizing coexistence mechanisms may operate at larger spatial scales in this community. Because foliar pathogens in this grassland community are multi-host and have small fitness impacts, they are unlikely to promote negative frequency-dependence or plant species coexistence in this system. Still, given that pathogen community composition differentiates across host species, some more subtle feedbacks between host relative abundance and pathogen community composition, damage, and fitness impacts are possible, which could in turn promote either coexistence or competitive exclusion.

## Introduction

Pathogens are ubiquitous in ecological communities (Burdon 1993, Gilbert 2002, Lafferty et al. 2008). Because they affect host demographic rates, pathogens are often expected to regulate host species population growth (Burdon 1982, Burdon and Chilvers 1982). Most pathogens infect only a subset of the available host species, so their incidence, and by extension their impacts, may be host-specific (Gilbert and Webb 2007, Beckstead et al. 2014, Parker et al. 2015). Host-specific population regulation can promote species coexistence by suppressing species when they become common and providing a relative advantage to rare species (Fig. 1). This pathogen-mediated negative frequency-dependence, sometimes called the Janzen-Connell hypothesis (Janzen 1970, Connell 1971), has growing support in diverse plant communities, including tropical forests (Augspurger 1983, Augspurger and Kelly 1984, Gilbert 2005, Bagchi et al. 2010, Bever et al. 2015), temperate forests (Packer and Clay 2000), and temperate grasslands (Petermann et al. 2008).

**Figure 1.**
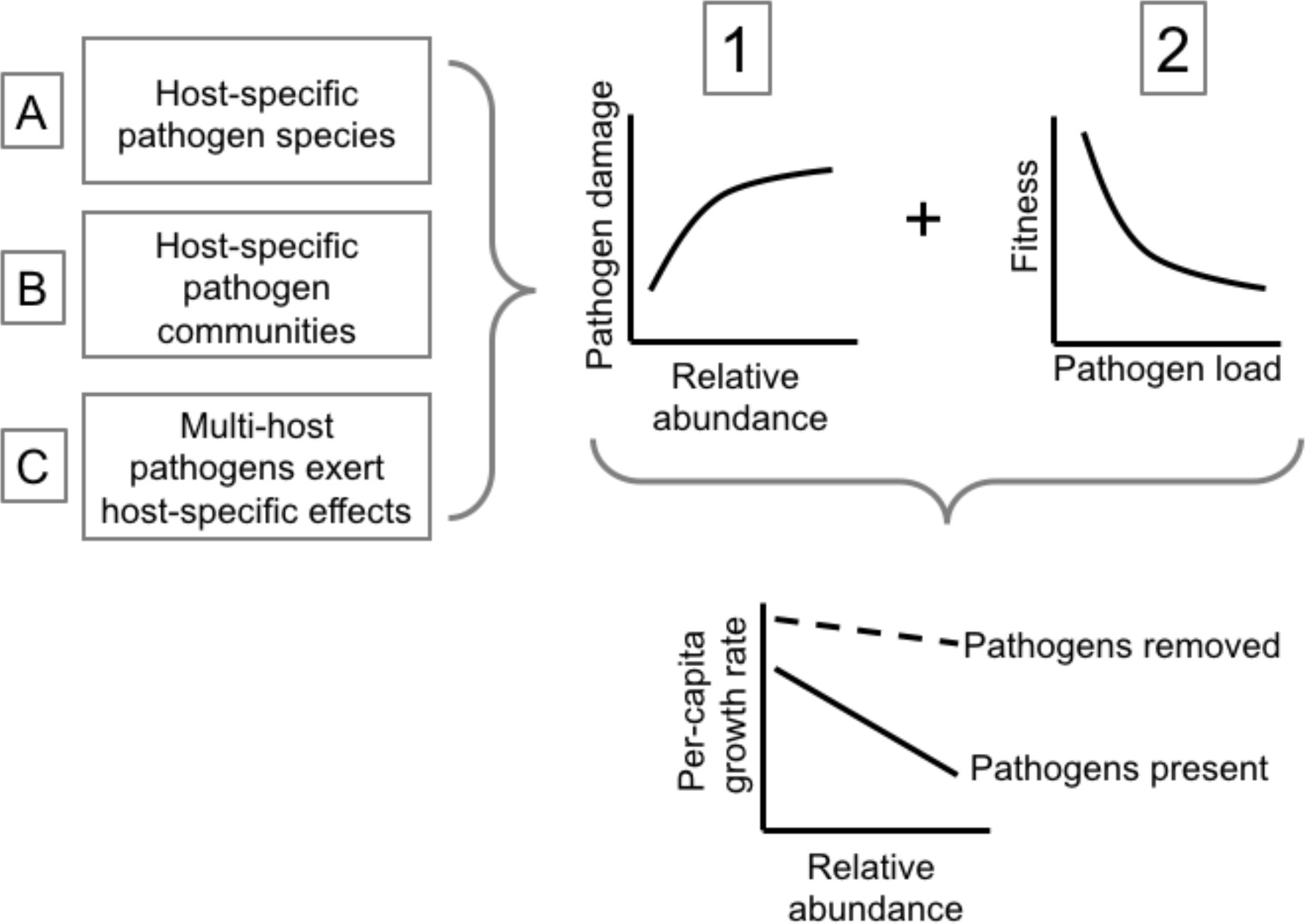
Pathogen-mediated frequency-dependence requires that: (1) pathogen damage increase with host species relative abundance, via mechanisms A-C, and (2) pathogen damage reduces fitness. The resulting decline in per-capita population growth rates with relative abundance can stabilize plant species coexistence.

At the same time, fungal pathogens of plants in natural systems often infect multiple hosts (Gilbert and Webb 2007, Kluger et al. 2008, Hersh et al. 2012, Spear 2017). In diverse plant communities, where the nearest neighbors may be heterospecifics, selection should favor multi-host pathogens (May 1991). Thus, we hypothesize that most plant pathogens are relatively host generalized (Spear et al. 2015, Spear 2017) and, by extension, that their attack rates and impacts do not respond to the relative abundance of a single host species. If this is the case, then, in contrast to the Janzen-Connell hypothesis, many pathogens may play little role in maintaining local host diversity and may even promote competitive exclusion or spatial turnover of species (Mordecai 2011, Spear et al. 2015). Few studies have assessed whether these alternatives to pathogen-mediated stabilization occur in nature (but see Mordecai 2013).

For pathogens to generate negative frequency-dependence and stabilize coexistence, they must: (1) disproportionately damage relatively common hosts, such that the amount or severity of damage depends on host relative abundance; and (2) reduce host population growth (Fig. 1). Condition (1) may occur either because (A) individual pathogen species are relatively specialized or exhibit host preference, (B) communities of multi-host pathogen species are differentially structured by plant species, or (C) multi-host pathogens exert host-specific impacts. In this paper, we measure frequency-dependent damage, host specificity, and impacts on population growth for foliar pathogens that infect co-occurring native and exotic grasses in a California grassland. Specifically, across six common grass species we: (i) quantified pathogen damage and linked it to plant species relative abundance; (ii) surveyed fungal community composition and pathogen sharing across host species; and (iii) measured the response of per-capita seed output to pathogen damage and plant relative abundance.

## Methods

### Study site & focal grass species

We conducted the study in grasslands in Jasper Ridge Biological Preserve (JRBP) at Stanford University, a 485-ha site in San Mateo County, CA (37°24’N, 122°13’30"W; 66 - 207 m), in 2015. JRBP has a Mediterranean climate, with cool (mean 9.2°C), wet winters and warm (mean 20.1°C), dry summers (total annual precipitation ~ 622.5 mm) (Ackerly et al. 2002). Plant growth begins with the onset of winter rains and plants senesce at the onset of summer.

We assessed the identities and impacts of foliar fungal pathogens on six common grass species: three exotic annuals, *Avena barbata, Bromus hordeaceus*, and *Bromus diandrus*; one exotic perennial, *Phalaris aquatica*; and two native perennials, *Stipa pulchra* and *Elymus glaucus*. To obtain a broader description of fungal diversity and host associations, we also isolated fungi from, but did not assess damage on, the common exotic grass species *A. fatua, Brachypodium distachyon*, and *Festuca perennis*. The exotic grasses were introduced to California in the mid-19^th^ century (Corbin and D’Antonio 2004).

We assessed the relationships between pathogen damage and host relative abundance (Condition 1), plant seed production and pathogen damage (Condition 2), and pathogen specialization and community composition (Conditions 1A-C).

### Impacts of plant species relative abundance on pathogen damage (Condition 1)

To measure pathogen burden across host species relative abundance (Condition 1), we visually measured the percentage of leaf area damaged by fungal pathogens (hereafter, pathogen damage) for the six focal grass species across 10 transects (yellow points in Fig. S1) that were established in areas where perennial species (either *S. pulchra*, *E. glaucus*, or *P. aquatica*) range from rare to common in a given 1-m^2^ plot; hereafter referred to as ‘perennial density transects’. As possible, we sampled multiple plants per transect and up to six haphazardly-selected leaves per plant, calculating average pathogen damage on each plant. To measure variation in pathogen damage within the growing season, we censused damage in these plots from March 11-16, 2015 (444 marked grass individuals) and from April 17-20, 2015 (163 of the marked grasses).

We tested whether pathogen damage correlated with host relative abundance in the plot (Condition 1) while controlling for other potential predictors: plant species, sampling month, and sampling structure (plot nested in transect as random effects), using normally-distributed errors (lmer function in the lme4 package; Bates et al. 2014). We assessed the most important predictors of pathogen damage by comparing the Akaike Information Criterion (AIC) values of models with all or a subset of the fixed effects of sampling month * species and species * frequency. We also modeled pathogen damage as a function of the dominant species at the plot scale as a categorical variable, but found no significant effect.

### Isolating foliar fungal pathogens (Conditions 1A-B)

To estimate fungal pathogen community composition and host ranges (Conditions 1A-B) at a preserve-wide scale, we cultured fungi from grasses along 24 transects that spanned a range of plant community composition, geographic location, and soil types (10 perennial density transects and 14 additional transects; Fig. S1 yellow and red points, respectively). Three of the pathogen survey transects ran through plots in the Jasper Ridge Global Change Experiment (GCE) (Zhu et al. 2016), where we sampled in ambient and water addition plots (water addition had no impact on fungal community composition). From March 19 – May 4, 2015, we collected and cultured from a symptomatic leaf from 772 grass individuals. We identified fungi (as described below) from 61 of the 444 marked plants from the damage survey and 219 additional plants. Sampling intensity varied among the nine grass species and transects based on availability of each species; the percentage of tissue pieces with growth varied among the hosts (Tables S1-S2). We excised tissue (< 2 mm^2^) from the advancing disease margin, surface-sterilized it in 70% EtOH followed by 10% household (Clorox) bleach (60 s each), then plated it on malt extract agar with 2% chloramphenicol (2% MEA). We pressed six of the segments onto 2% MEA to verify effectiveness of the surface sterilization; we observed no growth. We isolated morphologically distinct hyphae into pure culture on 2% MEA within 30 days. The Mordecai lab maintains reference strains (California Department of Food and Agriculture permit 3160).

### Identifying fungal species by DNA sequencing (Conditions 1A-B)

For each isolate, we extracted genomic DNA from fungal mycelium using REDExtract-N-Amp Tissue PCR Kit (Sigma-Aldrich, Inc.), following the manufacturer’s protocol. We amplified and sequenced the internal transcribed spacers (ITS) 1 and 2 and the 5.8S nuclear ribosomal gene using the primer pairs ITS-1F and ITS-4 (Gardes and Bruns 1993). For PCR amplification, we used a T100 thermal cycler (Bio-Rad Laboratories, Inc., Hercules, CA) and thermal cycling conditions following U’Ren et al. (2010). Following electrophoresis on a 1.5% agarose gel, we visualized PCR products using GelRed™ (Biotium Inc., Hayward, CA) and sent to them MCLAB (San Francisco, CA) for cleanup and bidirectional sequencing on an ABI 3730 XL sequencer.

We estimated the taxonomic relationships among the fungi in two ways. First, we assigned operational taxonomic units (OTUs) by manually editing all reads, automatically assembling bidirectional reads into consensus sequences using a minimum of 20% overlap and 85% sequence similarity, and clustering the 288 consensus sequences and two unidirectional sequences based on a minimum of 40% overlap and 90, 95, 97 and 99% sequence similarity using Sequencher (Gene Codes, Ann Arbor, MI). Second, because sequence similarity for the ITS region varies across fungal species (O’Brien et al. 2005), we built phylogenetic trees for groups of OTUs with similar sequences (1-52 isolates per dataset) (as described in Higginbotham et al. 2014, Spear 2017). Because not all sequences mapped onto named fungal species, we assigned each operational species a unique species code (Table S3). We treated isolates belonging to a species complex as a single species. All sequence data will be submitted to GenBank (accession numbers XXXX-XXXX).

### Analyses of fungal community composition and host associations (Conditions 1A-B)

To describe the fungal community (*N* = 290 isolates), we calculated (i) sampling efficacy using taxon accumulation curves, (ii) observed taxon richness, (iii) estimated lower bound of true richness, accounting for unseen taxa and correcting for under-sampling in highly diverse assemblages, with the iChao1 estimator (Chiu et al. 2014), (iv) rank-abundance distribution, and (v) diversity of taxa with Fisher’s alpha, which is robust to unequal sample sizes (Fisher et al. 1943, Magurran 2013) and the effective number of species (Jost 2006). We defined fungal species isolated ten or more times as abundant. We then contrasted fungal community composition across five grass species (Conditions 1A-B) using permutational multivariate analyses of variance (PERMANOVAs; Anderson 2001), with the adonis and pairwise.perm.manova functions (Oksanen et al. 2016, Hervé 2017). We visualized the differences using non-metric multidimensional scaling (NMDS). For these analyses (*N* = 99 isolates), we: (i) considered each grass species-by-perennial density transect combination to be a distinct community, excluding those communities with fewer than three isolates and *B. hordeaceus*, which only had one community with the three or more isolates; and (ii) created a matrix of pairwise community dissimilarities using the function vegdist with the Chao method, which is abundance-based and adjusted to consider unseen species (Chao et al. 2005, Oksanen et al. 2016). Finally, we made pairwise comparisons of the pathogen communities of the nine sampled grass species based on the observed number of shared species and using the Morisita-Horn index (*N* = 290 isolates, 200 bootstrap replicates).

To assess the specialization of the non-singleton fungal species (*N* = 228 isolates) (Condition 1A), we calculated the weighted specialization index d’, the degree to which a species deviates from random host associations, adjusted for the preserve-wide relative abundance of the hosts *A. barbata*, *B. diandrus*, *B. hordeaceus*, *S. pulchra*, and *E. glaucus* (see R code for data) (Blüthgen et al. 2006, Dormann et al. 2016). We classified the d’ values as low (0-0.33), moderate (0.34-0.67), and high (0.68-1) specialization (Blüthgen et al. 2006, Dormann et al. 2016).

All pathogen community analyses were conducted in *R* version 3.4.2, using the packages RVAideMemoire (Hervé 2017), vegan (Oksanen et al. 2016), SpadeR (Chao et al. 2016), bipartite (Dormann et al. 2016), fossil (Vavrek 2011), BiodiversityR (Kindt and Coe 2005), rich (Rossi 2011) and with custom commands (Gardener 2014).

### Pathogenicity tests (Conditions 1A-B)

We verified the pathogenicity of the fungal isolates by experimentally inoculating 99 isolates, representing 35 fungal species, onto the host species from which they were originally isolated (Table S4). In a greenhouse, we secured colonized or uncolonized (for paired controls) 2% ME agar plugs to healthy leaves using Parafilm (Sinclair and Dhingra 1995). We censused leaves for symptoms within one week, and compared the proportion of diseased leaves for each isolate to its paired control using bias-reduced generalized linear models (brglm function; Kosmidis 2013), with binomial errors and probit link functions (Table S4).

### Impacts of pathogen damage on per-capita seed production (Condition 1C, 2)

To assess the relationship between fecundity, community composition, and pathogen burden, we harvested seeds from 350 of the 444 grass individuals from which we had surveyed pathogen damage in the perennial density transects. We measured per-capita seed production, density of all host species, and pathogen damage for all focal species except *P. aquatica*, which occurs in monotypic stands with little variation in relative abundance. We standardized per-capita seed output and pathogen damage in either March or April by species using z-scores, and regressed pathogen damage on seed output (models with unstandardized variables produced similar results). We built separate models for pathogen damage in March versus April because they used the same individuals with a single seed output. Because variance in seed production differed across damage levels, we used quantile regression on the 25^th^, 50^th^, and 75^th^ percentiles of seed production (rq function, Koenker 2017). Finally, we linked fungal genera to pathogen damage in March and/or April (*N* = 65 isolates) and seed output (*N* = 56 isolates) (Condition 1C; we did not have enough samples to assess damage and seed output by fungal species). More comprehensively assessing fitness impacts would require measuring demographic rates on experimentally infected plants.

All pathogen damage and seed production statistical analyses were performed in *R* version 3.2.3 (R Development Core Team 2014) using the packages plyr (Wickham 2011), reshape (Wickham 2007), plotrix (Lemon 2006), ggplot2 (Wickham 2009), quantreg (Koenker 2017), lmerTest (Kuznetsova et al. 2016), lqmm (Geraci 2016), lme4 (Bates et al. 2014), piecewiseSEM (Lefcheck 2016), and lsmeans (Lenth and Love 2017).

## Results

### Pathogen damage across host species and relative abundance (Condition 1)

Focal species relative abundance was not significantly related to pathogen damage, counter to Condition 1 (Figs. 1 and 2). Instead, the strongest predictors of pathogen damage were host species and sampling month (Tables S5-S6). *A. barbata* and *B. hordeaceus* had the highest pathogen damage (Table S5). Pathogen damage was higher in April than in March (Fig. S2; paired two-tailed *t*-tests: mean difference = 0.0265, *t* = −4.1019, *df* = 162, *p* = 6.47 x 10^−5^).

**Figure 2.**
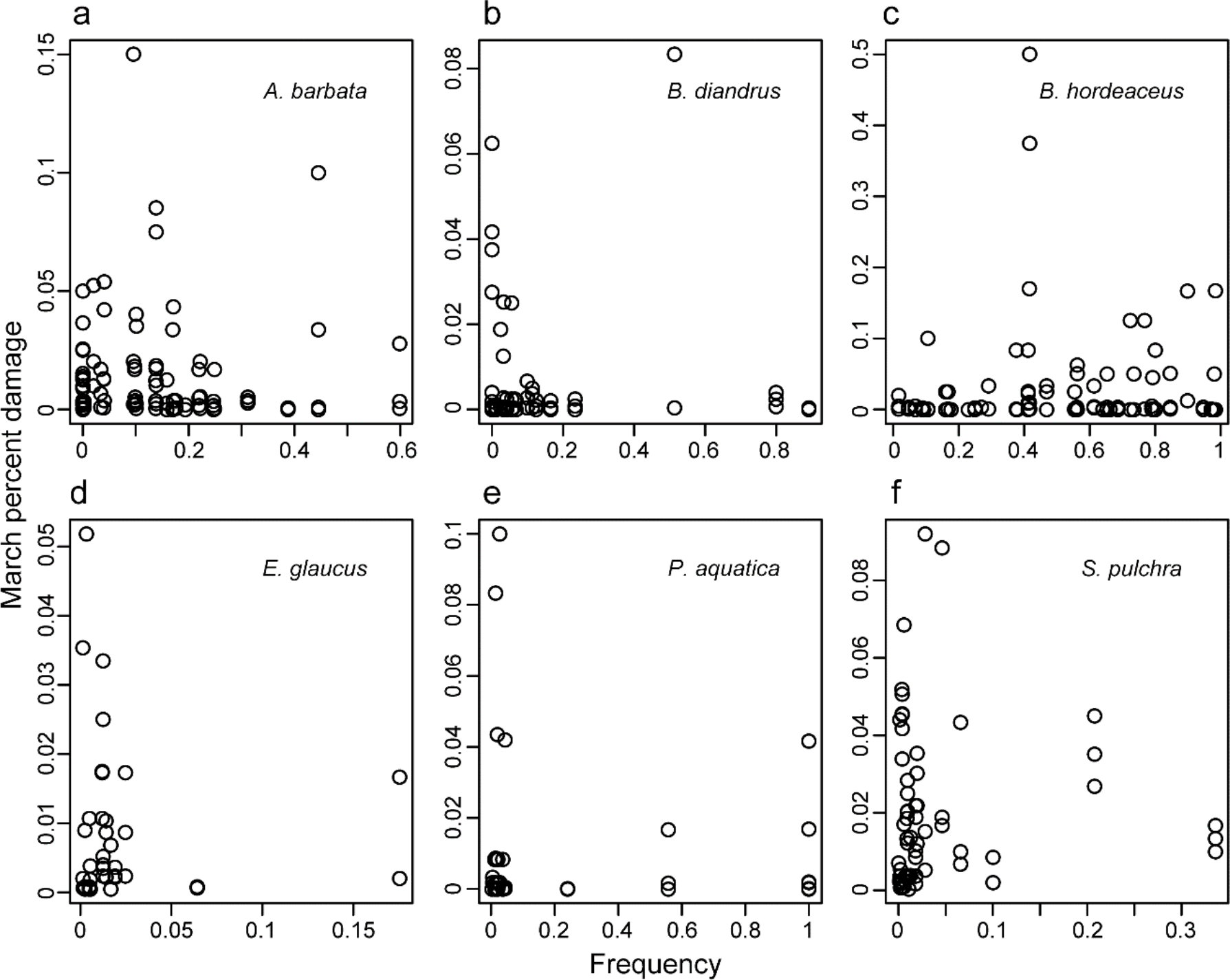
Relationship between focal species frequency in the plot and pathogen damage in March, for six focal species. Marginal R^2^ < 0.02 for all species.

### Fungal pathogen identities and diversity

We observed a diverse fungal community (Fig. S3). We cultivated 302 isolates (290 successfully sequenced) from 772 symptomatic leaves from nine plant species (Table S1). Considering 90 - 99% sequence similarity, the isolates represented 27 – 48 operational taxonomic units (OTUs), respectively (Fisher’s alpha index for diversity: 7.28 - 16.39). Hereafter, we designate all fungal species taxonomic affiliations based on the phylogenetic analyses. The fungal isolates represented 41 fungal species (iChao1 estimated species richness: 285.78, 95% CI = 85.91, 1375.24; Fisher’s alpha for diversity: 13.03, 95% CI = 8.91, 18.58; effective number of species = 18.29, 95% CI = 15.45, 21.14). Most fungal species were rare (56% were singletons or doubletons) and few were abundant (22% isolated >10 times) (Fig. S4). *Pyrenophora*, *Ramularia*, *Alternaria*, and *Parastagonospora* were the most common genera (Fig. 3 and Table S3). We experimentally confirmed the pathogenicity of 27% of the 99 isolates tested, representing eight of the nine common fungal species (Fig 3; Tables S3-S4).

**Figure 3.**
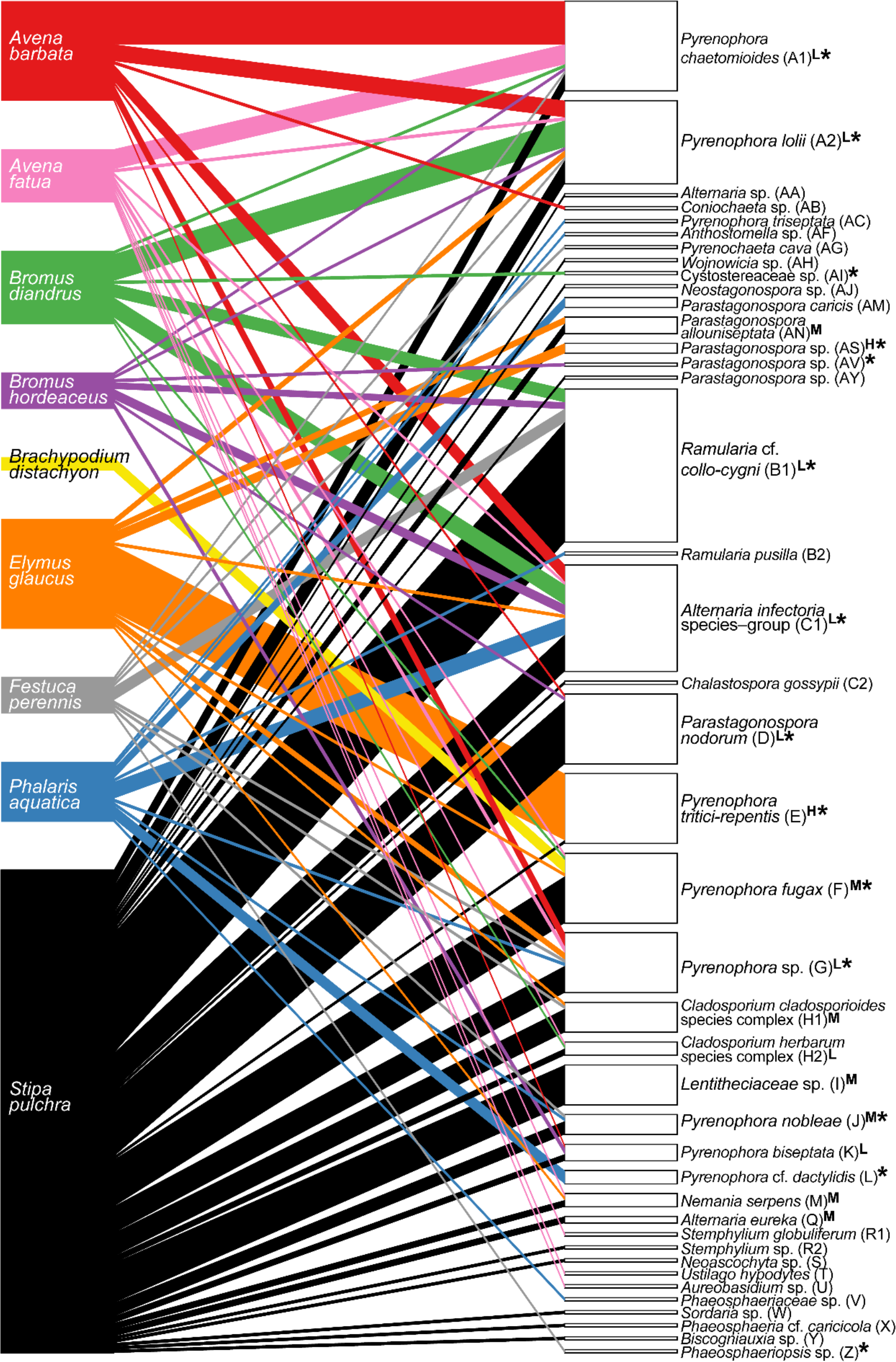
Bipartite network of 41 fungal pathogen species (right; *N* = 290 isolates) isolated from nine grass species (left), with widths proportional to isolation frequency. Fungal specificity (d’) is indicated for 17 common species (L = low, M = moderate, H = high). Experimentally confirmed pathogenicity is indicated with an asterisk.

### Fungal pathogen host associations and community similarity (Conditions 1A-B)

Most fungi were isolated from multiple hosts, counter to Condition 1A that fungal species are host-specific: 74% of the 19 non-singleton fungal species infected multiple hosts, averaging four host species (Fig. 3). Two fungi, *Pyrenophora lolii* (A2) and *Alternaria infectoria* species– group (C1), were isolated from seven of the nine grass species (Fig. 3). Concordantly, most non-singleton fungal species had low to moderate specificity (d’ median = 0.346; Table S7). An exception with high specificity (d’ = 0.924) was *Pyrenophora tritici-repentis* (E), which was isolated 20/21 times from *E. glaucus* and once from an *S. pulchra* individual in an *E. glaucus-*dominated plot; it comprised 61% of the 33 fungal isolates from *E. glaucus* leaves (Fig. 3; Table S7; Condition 1A). Two additional likely specialists were only isolated from the under-sampled grass species *P. aquatica* (*N* = 18 isolates): *Pyrenophora* cf. *dactylidis* (L) and *Parastagonospora caricis* (AM), comprising 39% of that host’s isolates (Fig. 3; Condition 1A).

The eight best-sampled grasses species shared one to eight fungal species with other grass species (median = 3; Table S8; Fig. 3). The average estimated pairwise similarity between host species was moderate (42%; min = 5% for *E. glaucus* – *P. aquatica*; max = 100% for *F. perennis* – *S. pulchra* and *A. barbata* – *A. fatua;* Table S8). By contrast, two grass species had low estimated fungal community similarity with other species: *E. glaucus* (5% – 11%), and *P. aquatica* (5% – 70%; Table S8; Fig. 3). Despite extensive pathogen sharing, the fungal pathogen communities of five of the focal grass species were significantly different, supporting Condition 1B that communities of shared pathogens are structured by host species (*N* = 99 isolates, PERMANOVA: *F*_4,15_ = 3.682, *R^2^* = 0.495, *p* = 0.001; Fig. 4).

**Figure 4.**
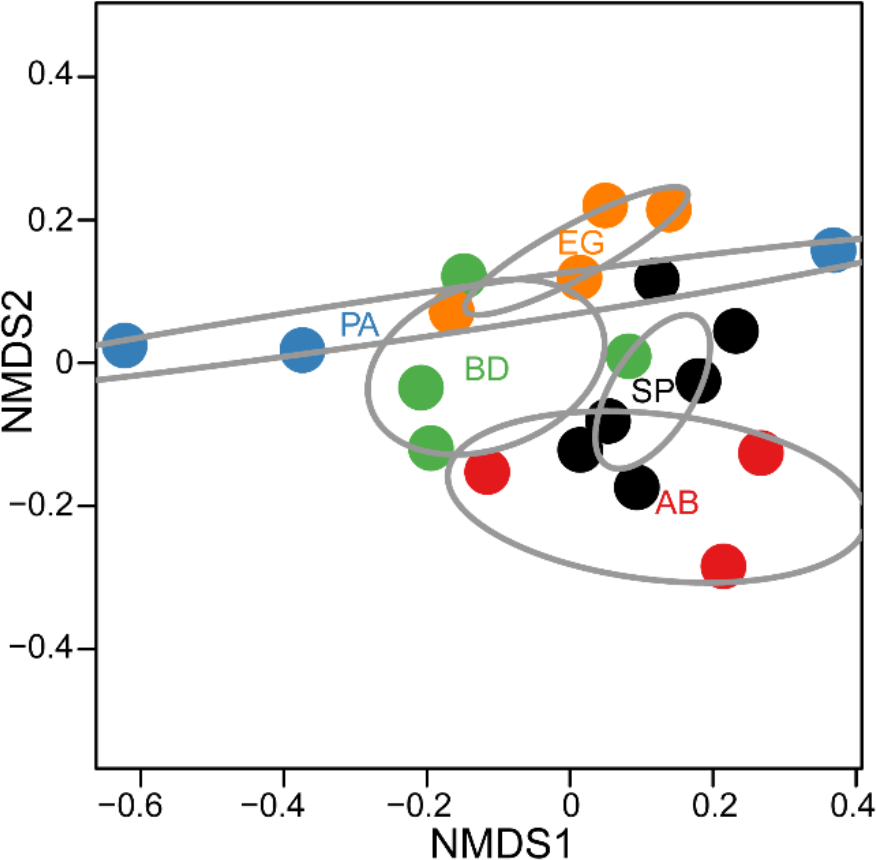
Dissimilarity of fungal pathogen community composition (*N* = 99 isolates, 21 fungal species) among five grass species (stress = 0.101). Data points represent fungal community composition across grass species-by-transect combinations, with 95% confidence ellipses around the centroid of each grass species. The fungal community of: (i) *S. pulchra* (SP) was significantly different from those of *A. barbata* (AB) (*p_adj_* = 0.037), *B. diandrus* (BD) (*p_adj_* = 0.037), *E. glaucus* (EG) (*p_adj_* = 0.04), and *P. aquatica* (PA) (*p_adj_* = 0.037); (ii) PA was significantly different from that of BD (*p_adj_* = 0.05); and (iii) BD was significantly different from that of AB (*p_adj_* = 0.05).

### Relationship between pathogen damage and per-capita seed output (Condition 2)

The relationships between pathogen damage and seed production were generally negative but highly variable, providing only weak support for Condition 2 that pathogens impact host fitness (Fig. 5a-b). Quantile regression showed that increasing pathogen damage in April was associated with reduced seed output at the 50^th^ percentile (i.e., for average-output individuals), but had no significant relationship for the 25^th^ or 75^th^ percentiles (i.e., for high- and low-output individuals) (Table S9; Fig. 5b). Contrary to expectations for negative frequency-dependence at local scales, focal species frequency had no association with either seed output (Fig. 5d) or pathogen damage (Fig. 2).

**Figure 5.**
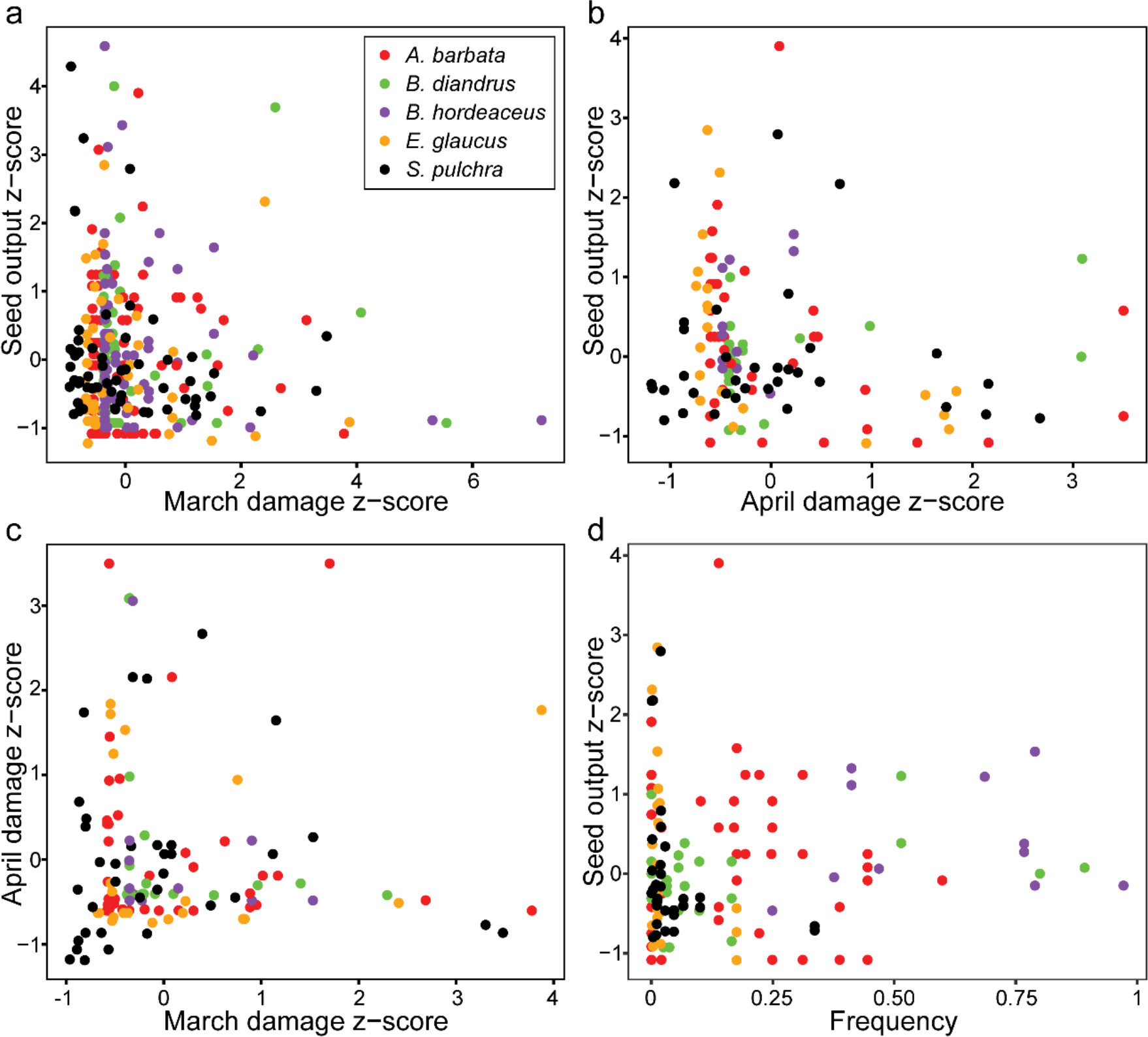
Pathogen damage in March versus seed production (a), pathogen damage in April versus seed production (b), pathogen damage in March versus April (c), and species frequency versus seed production (d). All values are expressed as z-scores calculated by species. Marginal R2 < 0.03 for all plotted relationships.

Particular fungal pathogen genera were not significantly associated with higher damage in April or lower seed production (Condition 1C), based on z-scores across host species (*N* = 65 individuals: 13 *A. barbata*, 8 *B. diandrus*, 17 *E. glaucus*, 22 *S. pulchra*, and 5 *P. aquatica*; Fig. S5). Small sample sizes precluded testing for host-specific impacts of all observed fungal species. Anecdotally, a single *S. pulchra* infected with *Sordaria* sp. (a singleton) had high pathogen damage and low seed production, consistent with a large pathogen impact.

## Discussion

### Limited potential for pathogen-mediated coexistence

Although pathogens are often hypothesized to promote plant species coexistence by generating negative frequency dependence (Gillett 1962, Augspurger 1983, Packer and Clay 2000, Petermann et al. 2008, Allan et al. 2010, Mangan et al. 2010, Bagchi et al. 2014, Bever et al. 2015, Whitaker et al. 2017), diverse natural communities should favor multi-host pathogens, decoupling pathogen abundance from the density of individual host species (May 1991, Spear et al. 2015). The degree of pathogen specialization and their role in promoting frequency dependence remains unresolved (Mordecai 2011). In a California grassland, we found ubiquitous pathogen damage: most plants and 57% of all surveyed leaves had pathogen damage. Yet the observed pathogen damage had no relationship with host relative abundance (contrary to Condition 1 for pathogens to cause negative frequency dependence; Fig. 1). Moreover, foliar pathogen damage was only weakly associated with lower seed production (Fig. 5a-b), providing limited support for pathogen impacts on plant fitness (Condition 2 in Fig. 1). While pathogens could also affect the outcome of competition by altering fitness differences between species, observational data (Fig. 5a-b) and preliminary experiments (Mordecai et al., unpublished) do not support strong, differential fitness impacts of foliar pathogen infection in this system.

The lack of a relationship between pathogen damage and host relative abundance (Fig. 2) is unsurprising given the extensive pathogen sharing among grass species (Table S8; Fig. 3; in contrast to Condition 1A in Fig. 1). Many pathogens are specific to a genus or family (Gilbert and Webb 2007, Barrett et al. 2009); thus, for the study grasses (all Poaceae), pathogen damage should be decoupled from the relative abundance of a single host species. It is also possible that pathogen damage responds to host relative abundance at the regional, rather than local, scale (Mitchell et al. 2002).

Despite pathogen sharing (Table S8; Fig. 3), fungal community composition subtly varied across grass species, partially supporting Condition 1B that pathogen communities differ across host species (Figs. 1, 3-4). In particular, the recently invading perennial grass *P. aquatica* (“Oakmead Herbarium: Arrivals, Weeds | Jasper Ridge Biological Preserve” 2018) had low pathogen damage and a distinct pathogen community, including two fungi with strong host affinities: *Pyrenophora cf. dactylidis* (L) and *Parastagonospora caricis* (AM). Moreover, several common fungi were relatively specialized (partially supporting Condition 1A [Fig. 1] that pathogens are host-specific), including *Pyrenophora tritici-repentis* (E) which was almost exclusively isolated from the native perennial *E. glaucus*. With limited sample sizes for each pathogen – host pair, we were unable to measure host-specific impacts of shared pathogens (Condition 1C) conclusively. In sum, although the multi-host fungal pathogens that dominate this California grassland community varied in their host affinities (Condition 1A), and pathogen communities were partially structured by host species (Condition 1B), neither mechanism was strong enough to generate frequency-dependent pathogen damage (Condition 1).

Our study focused on culturable fungi, which may be disproportionately host generalist, and on foliar pathogens, which may exert smaller fitness impacts than seedling damping off pathogens, root pathogens, and pathogens that castrate plants (Gilbert 2002). Those pathogens that impact different life stages and/or tissue types may shape the outcome of competition between plant species. A fuller assessment would require measuring fitness effects across life stages in experimental infections and incorporating them into population growth models.

The broad host ranges and limited fitness impacts of pathogens in this system contrast sharply with pathogen impacts on closely-related cultivated cereals such as barley, wheat, and oats. The fungal pathogen species we encountered are congeners of important cereal pathogens, including *Pyrenophora, Parastagonospora*, and *Ramularia* spp., which cause major yield losses and often specialize on host genotypes (Havis et al. 2015, McDonald and Stukenbrock 2016). Our results imply that naturally occurring host genetic and species diversity may mitigate the spread of highly virulent pathogens in wild grassland systems (McDonald and Stukenbrock 2016). Foliar pathogen load has declined with increased plant community diversity in natural grassland and old-field systems (Mitchell et al. 2002, Rottstock et al. 2014). Further, broad host ranges and minimal host impacts are well suited to pathogen persistence and spread in seasonal, mixed species grasslands like our California grassland site. Particularly in annual-dominated stands, pathogens must recolonize and spread during the limited winter and spring growing season each year, giving a selective advantage to multi-host pathogens.

### Conclusions

In one of the first studies to directly measure pathogen-mediated frequency-dependence and connect it to pathogen community composition, we showed that, contrary to prevailing hypotheses in other plant systems, foliar fungal pathogens are unlikely to promote plant coexistence in an invaded California grassland. Much of the evidence that pathogens maintain plant community diversity is based on spatial and temporal patterns of conspecific negative density-dependent mortality (e.g., Packer and Clay 2000, Klironomos 2002, Bell et al. 2006, Petermann et al. 2008, Bagchi et al. 2010, Mangan et al. 2010, Comita et al. 2010), with limited examination of fungal identity and host affinity (but see Parker and Gilbert 2007, Gilbert and Webb 2007, Hersh et al. 2012, Schweizer et al. 2013, Spear 2017). By contrast, our work suggests that pathogens have limited impacts on the outcome of competition when the burden is frequency-independent and fitness costs are minimal, which may commonly occur in natural systems.

## Acknowledgements

We appreciate help from Reuben Brandt, Joe Sertich, Johannah Farner, Fletcher Halliday, Bill Gomez, Meredith McClintock, Kabir Peay, Nora Dunkirk, Ryan Tabibi, Divya Ramani, Caroline Daws, Nona Chiariello, Philippe Cohen, Cary Tronson, and Steve Gomez, and funding from the Jasper Ridge Kennedy Endowment and the National Science Foundation (DEB-1518681).

## References

Ackerly, D., C. Knight, S. Weiss, K. Barton, and K. Starmer. 2002. Leaf size, specific leaf area and microhabitat distribution of chaparral woody plants: contrasting patterns in species level and community level analyses. Oecologia 130:449–457.

Allan, E., J. van Ruijven, and M. J. Crawley. 2010. Foliar fungal pathogens and grassland biodiversity. Ecology 91:2572–2582.

Augspurger, C. K. 1983. Seed dispersal of the tropical tree, Platypodium elegans, and the escape of its seedlings from fungal pathogens. The Journal of Ecology 71:759–771.

Augspurger, C. K., and C. K. Kelly. 1984. Pathogen mortality of tropical tree seedlings: experimental studies of the effects of dispersal distance, seedling density, and light conditions. Oecologia 61:211–217.

Bagchi, R., R. E. Gallery, S. Gripenberg, S. J. Gurr, L. Narayan, C. E. Addis, R. P. Freckleton, and O. T. Lewis. 2014. Pathogens and insect herbivores drive rainforest plant diversity and composition. Nature 506:85–88.

Bagchi, R., T. Swinfield, R. E. Gallery, O. T. Lewis, S. Gripenberg, L. Narayan, and R. P. Freckleton. 2010. Testing the Janzen-Connell mechanism: pathogens cause overcompensating density dependence in a tropical tree. Ecology Letters 13:1262–1269.

Barrett, L. G., J. M. Kniskern, N. Bodenhausen, W. Zhang, and J. Bergelson. 2009. Continua of specificity and virulence in plant host–pathogen interactions: causes and consequences. New Phytologist 183:513–529.

Bates, D., M. Maechler, B. Bolker, and S. Walker. 2014. lme4: Linear mixed-effects models using Eigen and S4.

Beckstead, J., S. E. Meyer, K. O. Reinhart, K. M. Bergen, S. R. Holden, and H. F. Boekweg. 2014. Factors affecting host range in a generalist seed pathogen of semi-arid shrublands. Plant Ecology 215:427–440.

Bell, T., R. P. Freckleton, and O. T. Lewis. 2006. Plant pathogens drive density-dependent seedling mortality in a tropical tree. Ecology Letters 9:569–574.

Bever, J. D., S. A. Mangan, and H. M. Alexander. 2015. Maintenance of plant species diversity by pathogens. Annual Review of Ecology, Evolution, and Systematics 46:305–325.

Blüthgen, N., F. Menzel, and N. Blüthgen. 2006. Measuring specialization in species interaction networks. BMC Ecology 6:9.

Burdon, J. 1993. The structure of pathogen populations in natural plant communities. Annual Review of Phytopathology 31:305–323.

Burdon, J., and G. Chilvers. 1982. Host density as a factor in plant disease ecology. Annual Review of Phytopathology 20:143–166.

Burdon, J. J. 1982. The effect of fungal pathogens on plant communities. The plant community as a working mechanism:99–112.

Chao, A., R. L. Chazdon, R. K. Colwell, and T. J. Shen. 2005. A new statistical approach for assessing similarity of species composition with incidence and abundance data. Ecology Letters 8:148–159.

Chao, A., K. H. Ma, T. C. Hsieh, and C.-H. Chiu. 2016. SpadeR: Species prediction and diversity estimation with R.

Chiu, C.-H., Y.-T. Wang, B. A. Walther, and A. Chao. 2014. An improved nonparametric lower bound of species richness via a modified good–turing frequency formula. Biometrics 70:671–682.

Comita, L. S., H. C. Muller-Landau, S. Aguilar, and S. P. Hubbell. 2010. Asymmetric Density Dependence Shapes Species Abundances in a Tropical Tree Community. Science 329:330–332.

Connell, J. H. 1971. On the role of natural enemies in preventing competitive exclusion in some marine animals and in rain forest trees. Dynamics of populations:298–312.

Corbin, J. D., and C. M. D’Antonio. 2004. Competition between native perennial and exotic annual grasses: Implications for an historical invasion. Ecology 85:1273–1283.

Dormann, C. F., J. Fruend, and B. Gruber. 2016. bipartite: Visualising Bipartite Networks and Calculating Some (Ecological) Indices.

Fisher, R. A., A. S. Corbet, and C. B. Williams. 1943. The Relation Between the Number of Species and the Number of Individuals in a Random Sample of an Animal Population. Journal of Animal Ecology 12:42–58.

Gardener, M. 2014. Community Ecology: Analytical Methods Using R and Excel. Pelagic Publishing Ltd.

Gardes, M., and T. D. Bruns. 1993. ITS primers with enhanced specificity for basidiomycetes - application to the identification of mycorrhizae and rusts. Molecular Ecology 2:113–118.

Geraci, M. 2016. lqmm: Linear Quantile Mixed Models.

Gilbert, G. S. 2002. Evolutionary ecology of plant diseases in natural ecosystems. Annual Review of Phytopathology 40:13–43.

Gilbert, G. S. 2005. Biotic interactions in the tropics: their role in the maintenance of species diversity. Pages 141–164 Dimensions of plant disease in tropical forests.

Gilbert, G. S., and C. O. Webb. 2007. Phylogenetic signal in plant pathogen–host range. Proceedings of the National Academy of Sciences 104:4979–4983.

Gillett, J. B. 1962. Pest pressure, an underestimated factor in evolution. Systematics Association Publication 4:37–46.

Havis, N. D., J. K. M. Brown, G. Clemente, P. Frei, M. Jedryczka, J. Kaczmarek, M. Kaczmarek, P. Matusinsky, G. R. D. McGrann, S. Pereyra, M. Piotrowska, H. Sghyer, A. Tellier, and M. Hess. 2015. Ramularia collo-cygni—An Emerging Pathogen of Barley Crops. Phytopathology 105:895–904.

Hersh, M. H., R. Vilgalys, and J. S. Clark. 2012. Evaluating the impacts of multiple generalist fungal pathogens on temperate tree seedling survival. Ecology 93:511–520.

Hervé, M. 2017. RVAideMemoire: Testing and Plotting Procedures for Biostatistics.

Higginbotham, S., W. R. Wong, R. G. Linington, C. Spadafora, L. Iturrado, and A. E. Arnold. 2014. Sloth Hair as a Novel Source of Fungi with Potent Anti-Parasitic, Anti-Cancer and Anti-Bacterial Bioactivity. PLOS ONE 9:e84549.

Janzen, D. H. 1970. Herbivores and the number of tree species in tropical forests. The American Naturalist 104:501–528.

Jost, L. 2006. Entropy and diversity. Oikos 113:363–375.

Kindt, R., and R. Coe. 2005. BiodiversityR. Tree diversity analysis. A manual and software for common statistical methods for ecological and biodiversity studies. World Agroforestry Centre (ICRAF), Nairobi.

Klironomos, J. N. 2002. Feedback with soil biota contributes to plant rarity and invasiveness in communities. Nature 417:67–70.

Kluger, C. G., J. W. Dalling, R. E. Gallery, E. Sanchez, C. Weeks-Galindo, and A. E. Arnold. 2008. Host generalists dominate fungal communities associated with seeds of four neotropical pioneer species. Journal of Tropical Ecology 24:351–354.

Koenker, R. 2017. quantreg: Quantile Regression. R, Vienna.

Kosmidis, I. 2013. brglm: Bias reduction in binomial-response Generalized Linear Models.

Kuznetsova, A., P. B. Brockhoff, and R. H. B. Christensen. 2016. lmerTest: Tests in Linear Mixed Effects Models.

Lafferty, K. D., S. Allesina, M. Arim, C. J. Briggs, G. De Leo, A. P. Dobson, J. A. Dunne, P. T. J. Johnson, A. M. Kuris, D. J. Marcogliese, N. D. Martinez, J. Memmott, P. A. Marquet, J. P. McLaughlin, E. A. Mordecai, M. Pascual, R. Poulin, and D. W. Thieltges. 2008. Parasites in food webs: the ultimate missing links. Ecology Letters 11:533–546.

Lefcheck, J. 2016. piecewiseSEM: Piecewise Structural Equation Modeling.

Lemon, J. 2006. Plotrix: a package in the red light district of R. R-News 6:8–12.

Lenth, R., and J. Love. 2017. lsmeans: Least-Squares Means.

Magurran, A. E. 2013. Measuring Biological Diversity. John Wiley & Sons.

Mangan, S. A., S. A. Schnitzer, E. A. Herre, K. M. L. Mack, M. C. Valencia, E. I. Sanchez, and J. D. Bever. 2010. Negative plant-soil feedback predicts tree-species relative abundance in a tropical forest. Nature 466:752–755.

May, R. M. 1991. A fondness for fungi. Nature 352:475–476.

McDonald, B. A., and E. H. Stukenbrock. 2016. Rapid emergence of pathogens in agroecosystems: global threats to agricultural sustainability and food security. Phil. Trans. R. Soc. B 371:20160026.

Mitchell, C. E., D. Tilman, and J. V. Groth. 2002. Effects of grassland plant species diversity, abundance, and composition on foliar fungal disease. Ecology 83:1713–1726.

Mordecai, E. A. 2011. Pathogen impacts on plant communities: unifying theory, concepts, and empirical work. Ecological Monographs 81:429–441.

Mordecai, E. A. 2013. Despite spillover, a shared pathogen promotes native plant persistence in a cheatgrass-invaded grassland. Ecology 94:2744–2753.

Oakmead Herbarium: Arrivals, Weeds | Jasper Ridge Biological Preserve. 2018, March 5. http://jrbp.stanford.edu/content/oakmead-herbarium-arrivals-weeds#Arrival.

O’Brien, H. E., J. L. Parrent, J. A. Jackson, J.-M. Moncalvo, and R. Vilgalys. 2005. Fungal Community Analysis by Large-Scale Sequencing of Environmental Samples. Applied and Environmental Microbiology 71:5544–5550.

Oksanen, J., R. Kindt, P. Legendre, and B. O’Hara. 2016. vegan: Community Ecology Package.

Packer, A., and K. Clay. 2000. Soil pathogens and spatial patterns of seedling mortality in a temperate tree. Nature 404:278–281.

Parker, I. M., and G. S. Gilbert. 2007. When there is no escape: the effects of natural enemies on native, invasive, and noninvasive plants. Ecology 88:1210–1224.

Parker, I. M., M. Saunders, M. Bontrager, A. P. Weitz, R. Hendricks, R. Magarey, K. Suiter, and G. S. Gilbert. 2015. Phylogenetic structure and host abundance drive disease pressure in communities. Nature 520:542–544.

Petermann, J. S., A. J. F. Fergus, L. A. Turnbull, and B. Schmid. 2008. Janzen-Connell effects are widespread and strong enough to maintain diversity in grasslands. Ecology 89:2399–2406.

R Development Core Team. 2014. R: A Language and Environment for Statistical Computing. R Foundation for Statistical Computing, Vienna, Austria.

Rossi, J.-P. 2011. rich: an R package to analyse species richness. Diversity 3:112–120.

Rottstock, T., J. Joshi, V. Kummer, and M. Fischer. 2014. Higher plant diversity promotes higher diversity of fungal pathogens, while it decreases pathogen infection per plant. Ecology 95:1907–1917.

Schweizer, D., G. S. Gilbert, and K. D. Holl. 2013. Phylogenetic ecology applied to enrichment planting of tropical native tree species. Forest Ecology and Management 297:57–66.

Sinclair, J. B., and O. D. Dhingra. 1995. Basic Plant Pathology Methods. CRC Press, Boca Raton, FL.

Spear, E. R. 2017. Phylogenetic relationships and spatial distributions of putative fungal pathogens of seedlings across a rainfall gradient in Panama. Fungal Ecology 26:65–73.

Spear, E. R., P. D. Coley, and T. A. Kursar. 2015. Do pathogens limit the distributions of tropical trees across a rainfall gradient? Journal of Ecology 103:165–174.

U’Ren, J. M., F. Lutzoni, J. Miadlikowska, and A. E. Arnold. 2010. Community Analysis Reveals Close Affinities Between Endophytic and Endolichenic Fungi in Mosses and Lichens. Microbial Ecology 60:340–353.

Vavrek, M. J. 2011. fossil: palaeoecological and palaeogeographical analysis tools. Palaeontologia Electronica 14:1T.

Whitaker, B. K., J. T. Bauer, J. D. Bever, and K. Clay. 2017. Negative plant-phyllosphere feedbacks in native Asteraceae hosts – a novel extension of the plant-soil feedback framework. Ecology Letters in press:n/a-n/a.

Wickham, H. 2007. Reshaping data with the reshape package. Journal of Statistical Software 21.

Wickham, H. 2009. ggplot2: Elegant Graphics for Data Analysis. Springer-Verlag, New York.

Wickham, H. 2011. The split-apply-combine strategy for data analysis. Journal of Statistical Software 40:1–29.

Zhu, K., N. R. Chiariello, T. Tobeck, T. Fukami, and C. B. Field. 2016. Nonlinear, interacting responses to climate limit grassland production under global change. Proceedings of the National Academy of Sciences 113:10589–10594.

